# Activity of human-specific Interlaminar Astrocytes in a Chimeric Mouse Model of Fragile X Syndrome

**DOI:** 10.1101/2025.02.26.640426

**Authors:** Alexandria Anding, Baiyan Ren, Ragunathan Padmashri, Maria Burkovetskaya, Anna Dunaevsky

## Abstract

Astrocytes, a subtype of glial cells, have multiple roles in regulating neuronal development and homeostasis. In addition to the typical mammalian astrocytes, in the primate cortex interlaminar astrocytes are located in the superficial layer and project long processes traversing multiple layers of the cerebral cortex. Previously, we described a human stem cell based chimeric mouse model where interlaminar astrocytes develop. Here, we utilized this model to study the calcium signaling properties of interlaminar astrocytes. To determine how interlaminar astrocytes could contribute to neurodevelopmental disorders, we generated a chimeric mouse model for Fragile X syndrome. We report that FXS interlaminar astrocytes exhibit hyperexcitable calcium signaling and are associated with dendritic spines with increased turnover rate.

## 1. Introduction

Astrocytes, a subtype of glial cells in the central nervous system (CNS), are crucial for maintaining CNS homeostasis [1]. They are intricately involved in neural circuits and influence various aspects of CNS function, including the regulation of neuronal development, synaptic transmission, ionic homeostasis, neurotransmitter recycling, metabolic regulation, and the maintenance of the blood-brain barrier [2–4]. Astrocyte excitability is manifested as elevations of cytosolic Ca^2+^. Astrocyte Ca^2+^ elevations can occur spontaneously as intrinsic oscillations in the absence of neuronal activity, can be triggered by neurotransmitters released during synaptic activity and can occur in response to sensory stimulation [5]. Increasing evidence highlights the heterogeneity of astrocyte functions, which arise from diverse morphologies and transcriptomic profiles across different brain regions [6]. However, much of this knowledge comes from studies on rodent astrocytes, which differ from their human counterparts in several key aspects [7–9], leaving the structural and functional properties of human astrocytes largely unexplored.

In comparison to rodents, human astrocytes exhibit distinct genomic profiles, larger territories, more complex morphologies, faster intracellular Ca^2+^ signaling, and morphological features unique to primates [8–10]. Astrocyte subtypes are regionally distributed within the brain, including protoplasmic astrocytes in the gray matter, fibrous astrocytes in the white matter, and other types such as velate astrocytes, perivascular astrocytes, Müller cells, and Bergmann glia [1]. Additionally, higher-order primates and humans possess astrocyte subtypes like interlaminar and varicose projection astrocytes, which are not found in rodents [1, 8, 11–13]. In the human cortex, interlaminar astrocytes (ILAs) reside in layer I and extend long processes that traverse multiple cortical layers, terminating in layers III/IV [12, 14]. While rudimentary interlaminar astrocytes can also be found in the rodent cortex, their processes are shorter and do not extend beyond layer I [12]. The functional roles of primate-specific interlaminar astrocytes in the development, maintenance, and function of cortical neural circuits remain poorly understood.

We recently developed a chimeric mouse model with human-induced pluripotent stem cell (hiPSC)-derived astrocytes, in which interlaminar astrocytes (ILAs) are present in the mouse cerebral cortex [15]. This model offers the first opportunity to investigate the functional properties of ILAs. In this study, we conducted Ca²⁺ imaging of ILA processes both in brain slices and *in vivo*, assessing their responses to canonical neurotransmitters. To assess a potential contribution of ILAs to neurodevelopmental disorders, we also engrafted astrocytes derived from hiPSCs of individuals with Fragile X syndrome (FXS), the most common inherited form of autism spectrum disorder. Given our previous findings that cultured astrocytes derived from FXS hiPSCs exhibit hyperexcitable Ca²⁺ signaling [16], we compared the Ca²⁺ signaling between FXS and control ILAs. Additionally, using multiphoton imaging of dendritic spines, we observed that dendrites near FXS ILAs have higher spine turnover rates compared to those near control ILAs, suggesting the FXS ILAs contribute to altered synaptic plasticity in FXS.

## 2. Results

### 2.1 Generation of hiPSCs-astrocyte chimeric mice

We differentiated human-induced pluripotent stem cells (hiPSCs) to neural progenitor cells (NPCs), and subsequently astrocytes [15, 17]. To visualize the hiPSC-astrocytes (hi-Astrocytes) in the chimeric mouse brain and to measure astrocyte Ca^2+^ signaling, we generated RFP or mScarlet and GCaMP6f-expressing astrocytes. We previously demonstrated that prior to engraftment, RFP-expressing hiPSC-astrocytes showed robust expression of canonical astrocyte markers [15]. Two-site injection resulted in the widespread distribution of hiPSC-astrocytes in the frontal cortex (Fig. 1A). While at what we presume to be sites of injection both ILA and deeper protoplasmic astrocytes are observed, further away, cells are mainly confined to layer 1 and likely comprised of both pial and subpial ILAs (Fig. 1B).

**Figure 1:**
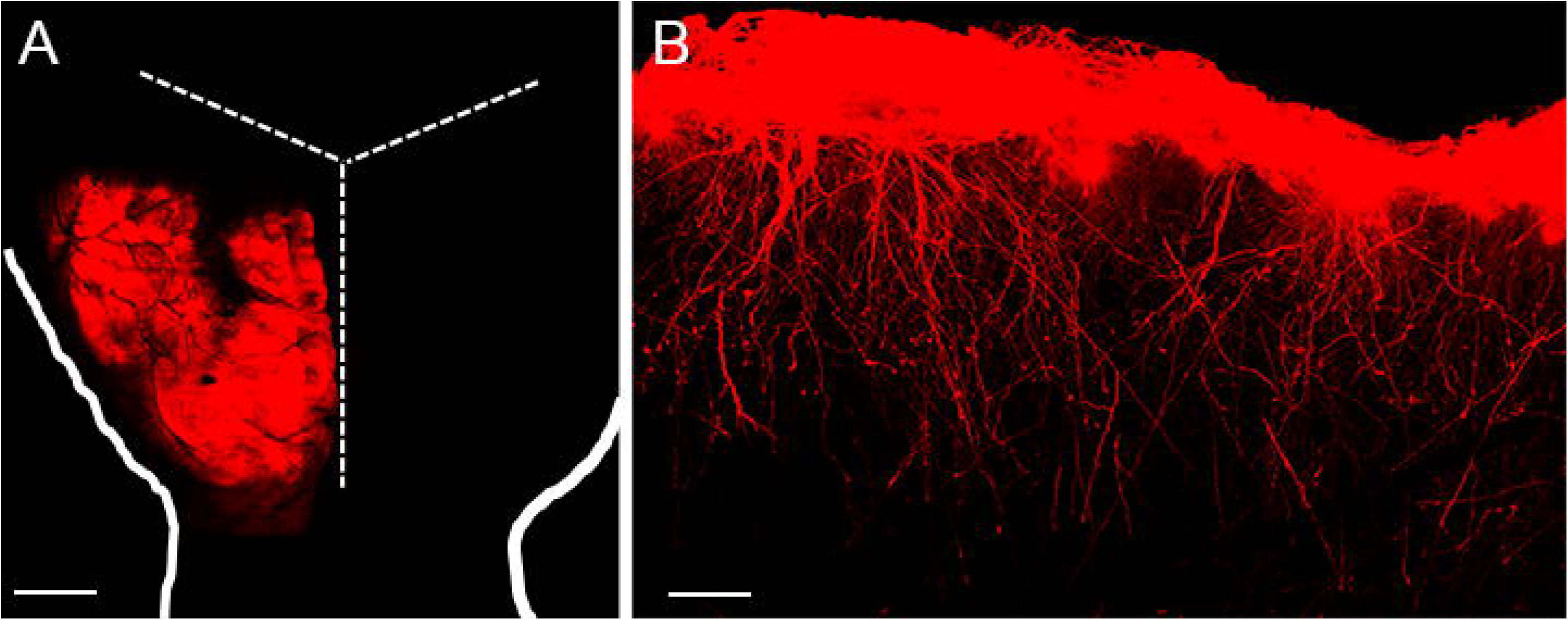
Chimeric mice engrafted with hiPSC-astrocytes develop interlaminar processes. **(A)** Whole mount view of a brain from a nine-month old mouse engrafted with RFP-expressing hiPSC-astrocytes. The dashed lines mark the skull sutures and the solid line outlines the brain. **(B)** Sagittal section from a 6-month engrafted mouse with RFP-expressing hiPSC-astrocytes. Interlaminar astrocytes are seen throughout the anterior-posterior (right-left) area. Scale Bar: 1mm in **A** and 200 microns in **B.**

### 2.2 Calcium signaling properties of interlaminar astrocytes

The dynamic Ca^2+^ properties of ILAs and their long processes has not been previously examined. We therefore imaged slices at 6 months post engraftment in cells that expressed GCaMP6f and the structural marker mScarlet. At this age Ca^2+^ signals in both cell bodies at the pial surface and the long ILA processes that project from the pial surface could be examined (Fig. 2A,B). To capture as much of the long ILA processes as possible, we performed volumetric 2-photon imaging of a 20-micron volume at 1 Hz and imaged both spontaneous and agonist-induced changes (Fig. 2). We observed very little spontaneous activity with none of the ILA somas exhibiting activity in the absence of an agonist. We next asked if hiPSC derived ILA cells are responsive to the canonical transmitters that have been examined extensively in mice; ATP (100 µM) to activate purinergic receptors, norepinephrine (NE, 50 µM) to activate noradrenergic receptors and carbachol (CA, 50 µM) to activate cholinergic receptors. We found that 56% and 44% of processes responded to ATP (N=141) and NE (N=100), respectively (Chi-Square, P=0.043). Interestingly, none of the ILAs responded to carbachol (data not shown). These data suggest that engrafted hi-Astrocytes that develop into ILAs express purinergic and adrenergic receptors and the molecular machinery for Ca^2+^ signaling. However, the lack of response to carbachol suggests low or absent expression of cholinergic receptors on ILAs.

**Figure 2:**
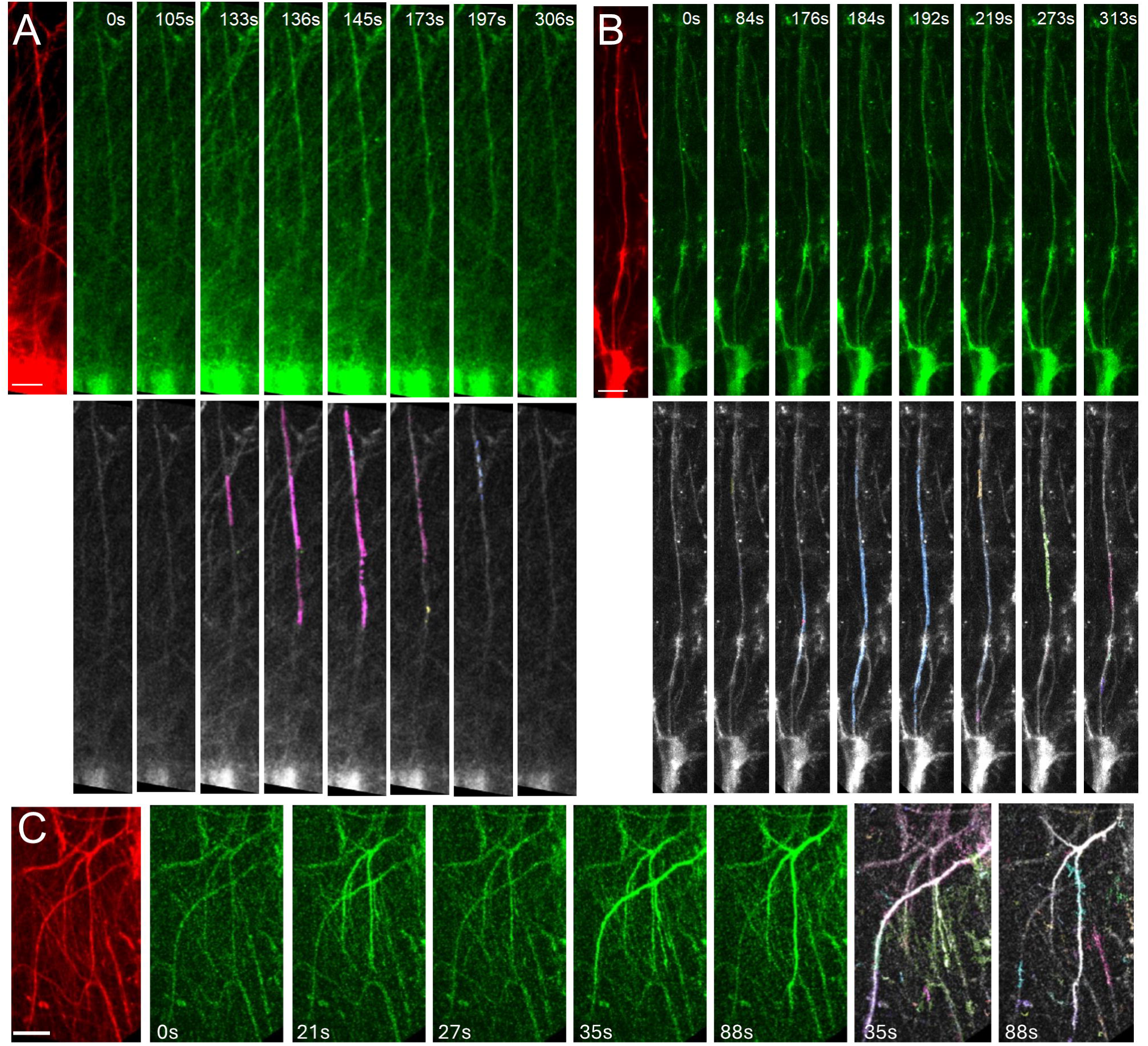
Imaging Ca^2+^ activity in the interlaminar astrocytes. Time-lapse imaging of ILA in a cortical slice from a 6-month-old chimeric mouse engrafted with immature astrocytes derived from control **(A)** or FXS **(B)** stem cells. ILAs expressing the structural marker mScarlet is shown (red). ATP-evoked Ca^2+^ activity in ILAs expressing GCaMP6f (green) with the AQuA-detected events in corresponding time frames shown. Propagation of the Ca^2+^ signal along the ILA process can be observed. **C.** In vivo imaging of ILA in an awake head restrained mouse. Shown here are mScarlet expressing ILA astrocytes, time-lapse imaging of Ca^2+^ signals in ILA processes and the AQuA-detected events. Scale bars: 25 µm.

We next characterized the properties of the Ca^2+^ signals in the somas and the processes. ILA somas, from engrafted control hi-Astrocytes, had robust Ca^2+^ responses to both ATP and NE that lasted for tens of seconds (Fig. 3A, Supplemental Data Movie 1). We did not observe a difference in the peak amplitude (dF/F), area under the curve or the duration of the ATP- and NE-induced responses in control ILA somas (ATP N=36 and NE N=27 somas, multiple Mann-Whitney tests P>0.05). Since ILAs don’t appear to have fine processes that are found on protoplasmic astrocytes, we only considered event sizes larger than 3 μm^2^ and found that following ATP or NE application Ca^2+^ events spanned a range of 3-105 μm^2^, however the distribution of ATP-induced Ca^2+^ events was shifted towards larger events compared to NE-induced events (Fig. 3B, Kolmogorov-Smirnov test, P=0.025). The maximum event size is constrained by the length of the outlined processes that are observed within the slices. However, the larger event areas with ATP were not due to longer regions of interest, as in fact they were longer in the NE treated slices (ATP: 91.99±4.28 µm (N=28); NE: 114±5.3 µm (n=31), unpaired t-test P=0.002). We next characterized the Ca^2+^ signaling properties based on event sizes dividing them into small (3-10 μm^2^) and large events (>10 μm^2^) (Fig. 3C). Despite the difference in the event areas, the dynamic properties of the ATP- and NE-induced Ca^2+^ events in control cells were not different in any of the parameters examined (Fig. 3C). These studies demonstrate that the long ILA processes exhibit Ca^2+^ signaling in response to canonical agonists.

**Figure 3:**
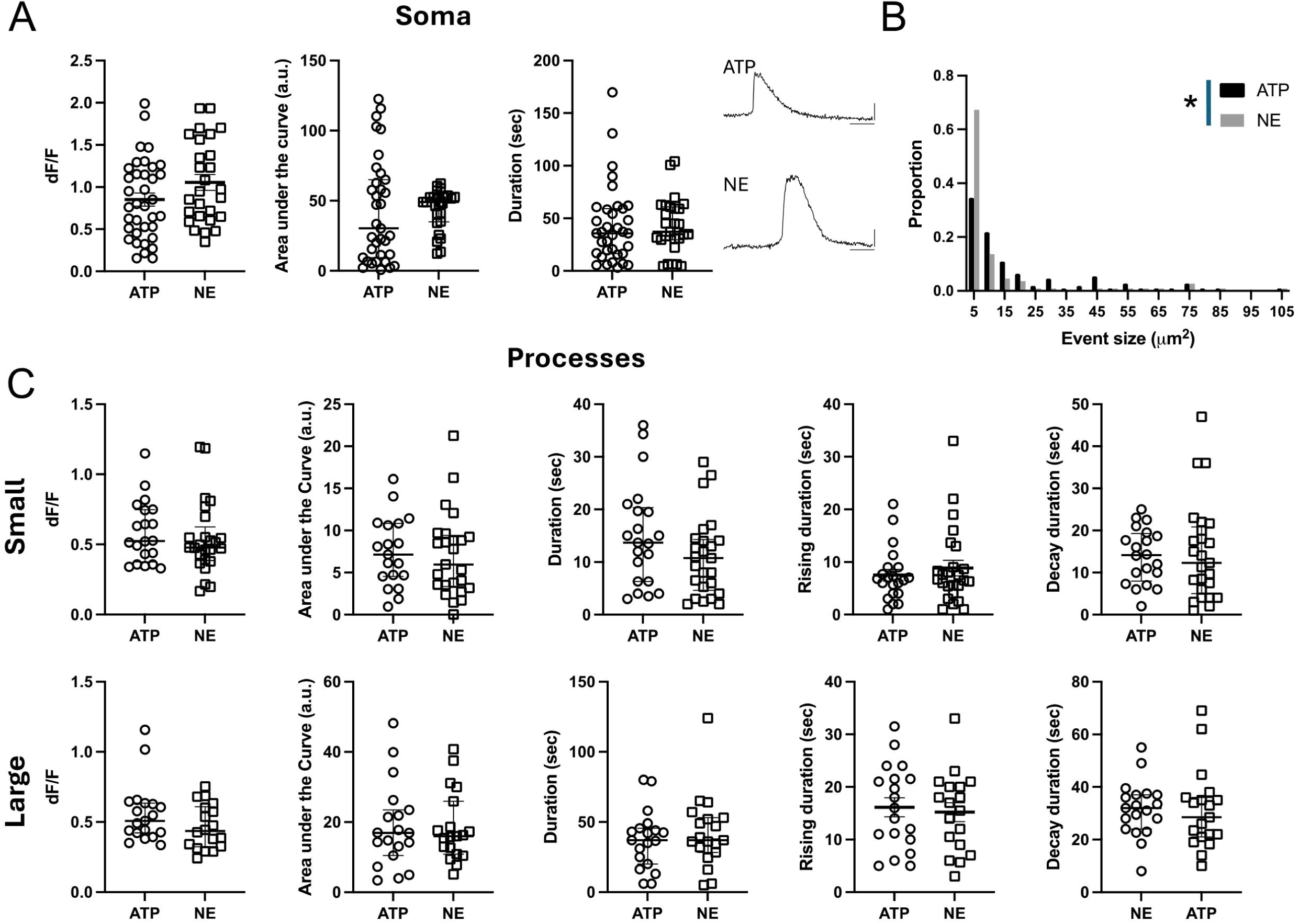
Ca^2+^ signaling properties in interlaminar astrocytes. **(A)** ATP and NE-evoked Ca^2+^ activity in ILA somas in cortical slices. Data is shown for Ca^2+^ event amplitude, area under the curve and duration. ATP: 6 mice, 13 slices, N = 36; NE: 4 mice, 8 slices, N = 27. Example traces are shown for ATP- and NE-evoked Ca^2+^ signals. **(B)** Frequency distribution for size of the Ca^2+^ events in ILA processes with ATP and NE application. Kolmogorov-Smirnov test, P=0.025. **C.** ATP and NE-evoked Ca^2+^ activity in ILA processes in cortical slices. Data is shown for Ca^2+^ event amplitude, area under the curve, duration, rise time to peak and decay time for small events (top panel) and large events (bottom panel). No significant differences were observed in Ca^2+^ signaling properties between ATP- and NE-evoked responses in the ILA processes. ATP: 7 mice, 17 slices, N = 18-20 processes; NE: 5 mice, 13 slices, N = 17-25 processes. Multiple Mann-Whitney tests. Mean ± SEM or median ± interquartile range are shown.

To determine if the Ca^2+^ signaling observed in ILA processes occur in the intact brain, we performed 2-photon in vivo imaging of hi-Astrocyte engrafted mice through a cranial window in 6-month old awake head-restrained conditions. Similarly to the observed Ca^2+^ activity in slices, we observed that long, thin processes in cortical Layer 1 exhibited robust Ca^2+^ responses in vivo (Fig. 2C). The activity is likely to be induced by the movement of the mouse (Supplemental Data Movie 2).

### 2.3 ILA process length is not significantly altered in FXS

Impairments in astrocyte function are increasingly associated with neurodevelopmental disorder [18]. However, how ILAs that are specific to humans and non-human primates are altered in neurodevelopmental disorders has not been examined before. We have utilized the human astrocyte mouse chimera model with astrocytes derived from FXS hiPSC or hESC with deletion of FMR1 to determine if ILA development and function is altered. We have previously described the gradual development of ILA processes in the engrafted mice with few processes observed at 3 months and full “palisade-like” processes observed at 9 months of age [15]. Here we first asked if process length was altered in FXS ILAs. Due to the large number of processes in the “palisade-like” distribution, we were unable to reliably track the processes of individual cells. Instead, we measured the distance of the “interlaminar palisade” from the pial surface in FX11-9u (CTR) and FX11-7 (FXS) hiPSC-astrocyte engrafted mice at 3 and 9 months of age (Fig. 4B). While the processes displayed a significant age factor (F (2,40)=16.55, P<0.0001) and FXS ILA processes tended to be longer, the genotype factor was not significant (F (1,40)=3.129, P=0.08) nor was there a significant interaction (F (2, 40) = 0.7415, P=0.48) (3 months: 145.00 ± 38.81 and 228.9 ± 29.90 μm, 9 months: 572.7 ± 81.49 and 673.4 ± 89.44 μm). Thus, we conclude that there was no gross alteration in the development of ILAs in FXS.

**Figure 4:**
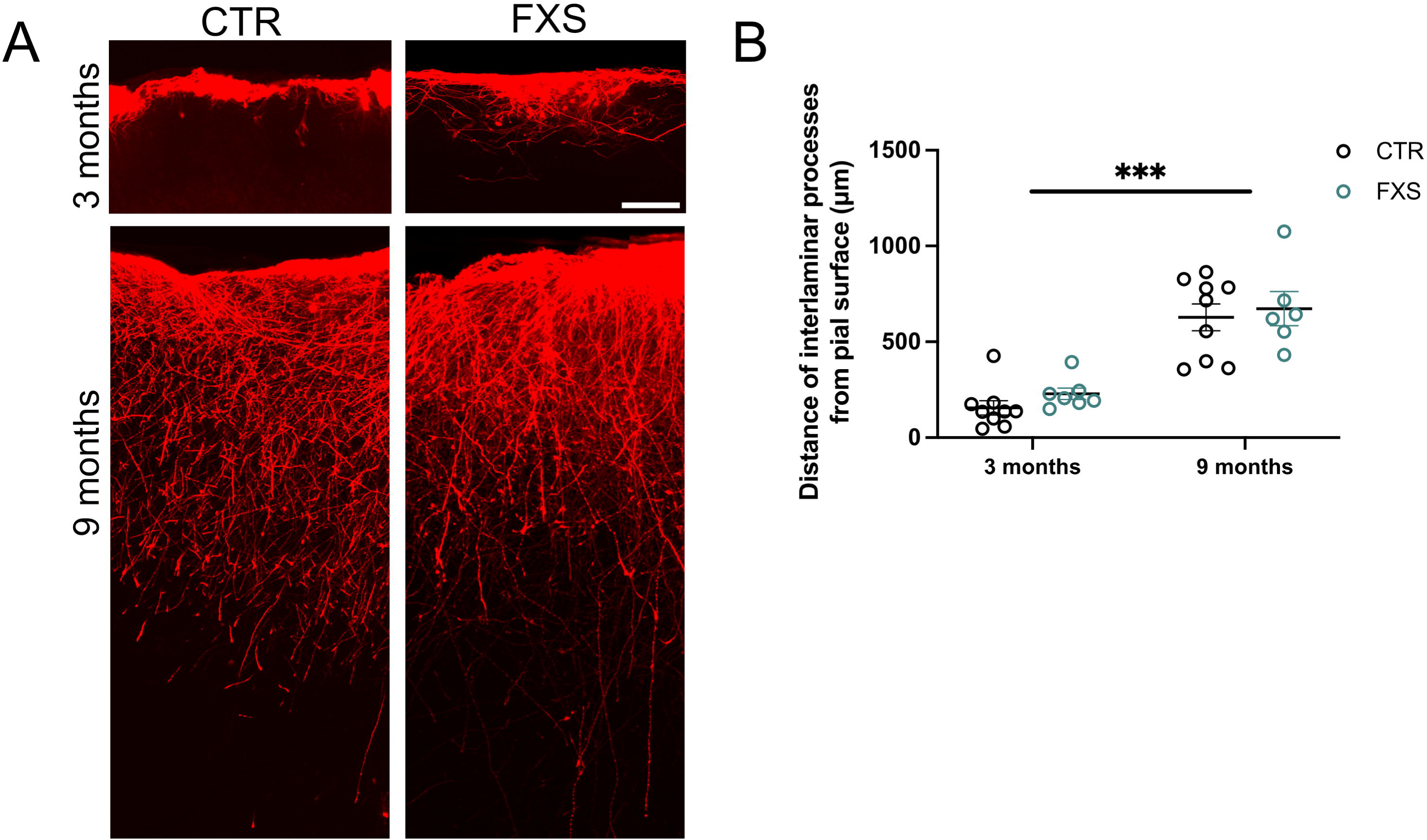
Interlaminar astrocyte process length is not altered in FXS. **(A)** Control and FXS human astrocytes expressing RFP in the cortex of 3 and 9-month-old chimeric mice. Scale bar, 100 µm. **(B)** Quantification of the distance traversed by CTR and FXS ILA processes in the 3 and 9-month-old chimeric mice. N = 5–9 sections from 2 to 3 chimeric mice per group. Two-way ANOVA. Age factor (F (2,40)=16.55, P<0.0001)

### 2.4 FXS ILAs have increased calcium signaling

We next compared the dynamic Ca^2+^ properties of engrafted astrocytes derived from control (3 lines, same data as in Figure 3) and FXS (2 lines) human stem cells. At 6 months of age we observed that the somas of FXS ILAs had approximately 60% and 100% increase in the duration of the response to ATP and NE, respectively (Fig. 5A and 6A, ATP, N=36 control and N=33 FXS somas, P=0.0002; NE, N=27 control and N=12 FXS somas, P=0.008. Multiple Mann-Whitney tests). We did not observe a difference between the peak amplitude (dF/F) of the soma responses to either ATP or NE between control and FXS cells (Fig. 5A and 6A). Interestingly, analysis of hiPSCS-astrocytes from a subset of lines after 4 months of engraftment, revealed multi-peak Ca^2+^ transients in most cells in response to bath-application of 100 µM ATP (Fig. S1A). Unlike the increased duration observed at 6 months, at 4 months we observed increased peak amplitude in engrafted FXS hiPSC-astrocytes (Fig. S1B, CTR: 0.77 ± 0.087 dF/F, FXS: 1.18 ±.065 dF/F, P = 0.0023, N = 6-8 slices from 4-5 mice in each group). These data demonstrate that FXS derived astrocytes that develop in vivo, exhibit hyperexcitability of Ca^2+^ in response to multiple agonists.

**Figure 5:**
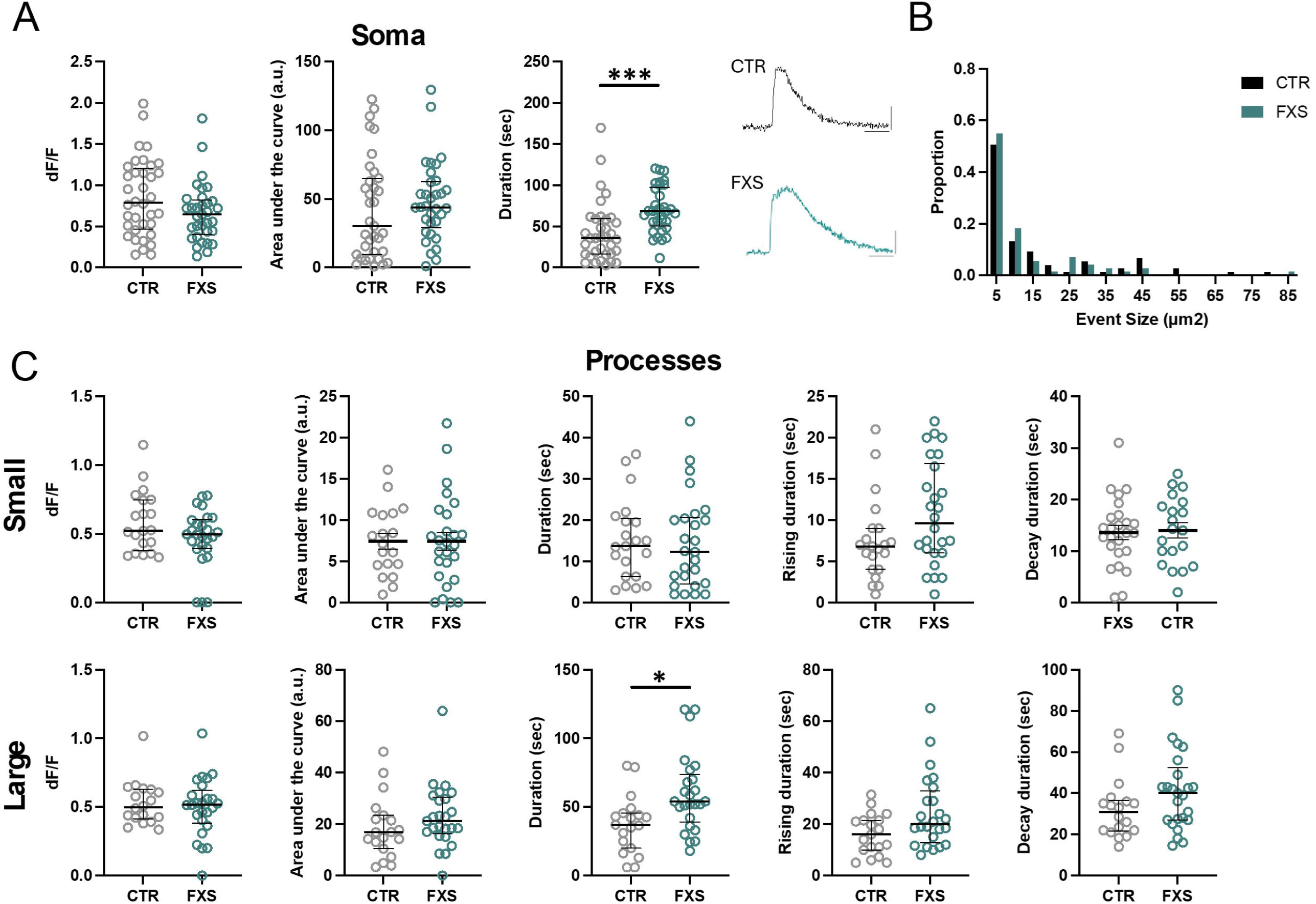
Enhanced ATP-evoked Ca^2+^ signaling in FXS interlaminar astrocytes. **(A)** ATP-evoked Ca^2+^ activity in soma of CTR and FXS ILAs in cortical slices. Data is shown for Ca^2+^ event amplitude, area under the curve and duration. FXS astrocyte soma exhibited increased Ca^2+^ event duration. CTR: 6 mice, 13 slices, N = 36 somas; FXS: 6 mice, 10 slices, N = 33 somas. Multiple Mann-Whitney tests, P=0.0002. Example traces of ATP-evoked Ca^2+^ signals in CTR and FXS ILA soma. **(B).** Frequency distribution for size of the Ca^2+^ events in processes in CTR and FXS astrocytes. Kolmogorov-Smirnov test, P>0.05. **(C)** ATP-evoked Ca^2+^ activity in ILA processes of CTR and FXS astrocytes in cortical slices. Data is shown for Ca^2+^ event amplitude, area under the curve, duration, rise time to peak and decay time for small events (top panel) and large events (bottom panel). A significant increase in the Ca^2+^ event duration for large events was observed in the FXS ILA processes. CTR: 7 mice, 17 slices, N = 18-20 processes; FXS: 9 mice, 16 slices, N = 24-26 processes for FXS, multiple Mann-Whitney tests P=0.029. Mean ± SEM and median ± interquartile range are shown.

**Figure 6:**
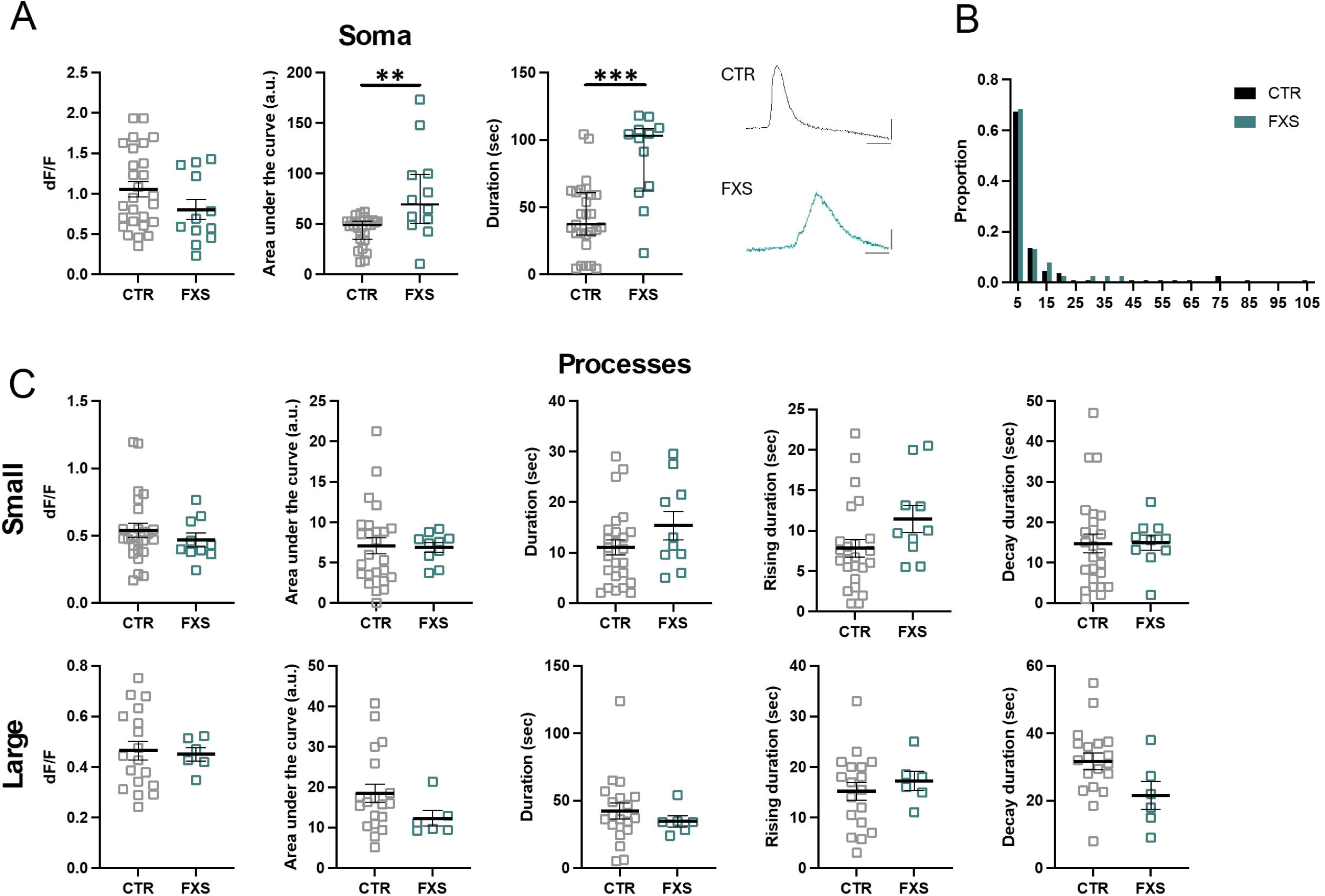
Enhanced NE-evoked Ca^2+^ signaling in FXS interlaminar astrocytes. **(A)** NE-evoked Ca^2+^ activity in somas of CTR and FXS ILAs in cortical slices. Data is shown for Ca^2+^ event amplitude, area under the curve and duration. FXS astrocytes exhibited increased Ca^2+^ event duration and area under the curve. CTR: 4 mice, 8 slices, N = 27 somas; FXS: 3 mice, 4 slices, N = 12 somas. Multiple Mann-Whitney tests, P=0.008 and P=0.006. Example traces of NE-evoked Ca^2+^ signals in CTR and FXS ILA soma. (**B)** Frequency distribution for size of the Ca^2+^ events in processes in CTR and FXS ILAs. Kolmogorov-Smirnov test, P>0.05. **(C)** NE-evoked Ca^2+^ activity in ILA processes of CTR and FXS astrocytes in cortical slices. Data is shown for Ca^2+^ event amplitude, area under the curve, duration, rise time to peak and decay time for small events (top panel) and large events (bottom panel). No significant changes were observed in the FXS astrocyte processes. CTR: 5 mice, 13 slices, N = 17-25 processes; FXS: 4 mice, 5 slices, N = 6-10 processes. Multiple Mann-Whitney tests.

We next examined if the Ca^2+^ hyperexcitability observed in FXS somas is also observed in processes. Analysis of the ILA processes revealed no difference in the size of the area of ATP- and NE -induced Ca^2+^ events detected by AQuA between CTR and FXS (Fig. 5B and 6B). However, analysis of small and large ILA events determined that while most parameters were similar between CTR and FXS (Fig. 5C and 6C), a 62% increase in the duration of large ILA events in response to ATP was observed in FXS (CTR: N=19, FXS=26 processes, P<0.05). Overall, these data indicate hyperactive calcium signaling in FXS ILAs.

### 2.5 Increased dendritic spine turnover in mice engrafted with FXS ILAs

Cumulative evidence showed that astrocytes regulate spine formation and elimination to maintain dynamic and precise neuronal connections. Previous studies identified increased spine dynamics in the *Fmr1* KO mice [19–21]. Several studies described the contribution of astrocytes to the neuronal abnormalities in the mouse model of FXS [22–25]. Thus, we asked whether human astrocytes contribute to the altered spine plasticity in FXS by repeatedly imaging the same neuronal dendritic segments in vivo in the chimeric mice. Excitatory neurons in the chimeric mice were labeled with eGFP through viral injection [26]. Dendritic spines in proximity of RFP expressing hiPSC-astrocytes (distance < 20 µm) were imaged through the cranial window twice with 4-day interval (Fig. 7A). Dendritic spine density on dendrites in vicinity of FXS hiPSC-astrocytes showed a trend towards increased density (Fig. 7B, CTR: 2.54± 0.21, FXS: 3.48 ± 0.42, P = 0.082). Though there was only a trend towards an increased percentage of newly formed dendritic spines in the FXS group compared with that in the CTR group (Fig. 7C, CTR: 20.89 ± 5.78%, FXS: 34.86 ± 6.34 %, P = 0.16, 16-28 regions from N=5 mice in each group), the percentage of spine elimination on dendrites in FXS group was higher than that in CTR group (Fig. 7D, CTR: 19.11 ± 199 %, FXS: 29.1 ± 1.33 %, P = 0.003) as was the turnover rate (TOR) [20, 27] (Fig. 7E, CTR: 0.21 ± 0.04, FXS: 0.32 ± 0.03, P = 0.037). These results demonstrate an increase in host dendritic spine plasticity attributed to engrafted FXS human astrocytes, indicating the role of astrocytes in the spine dynamics in FXS.

**Figure 7:**
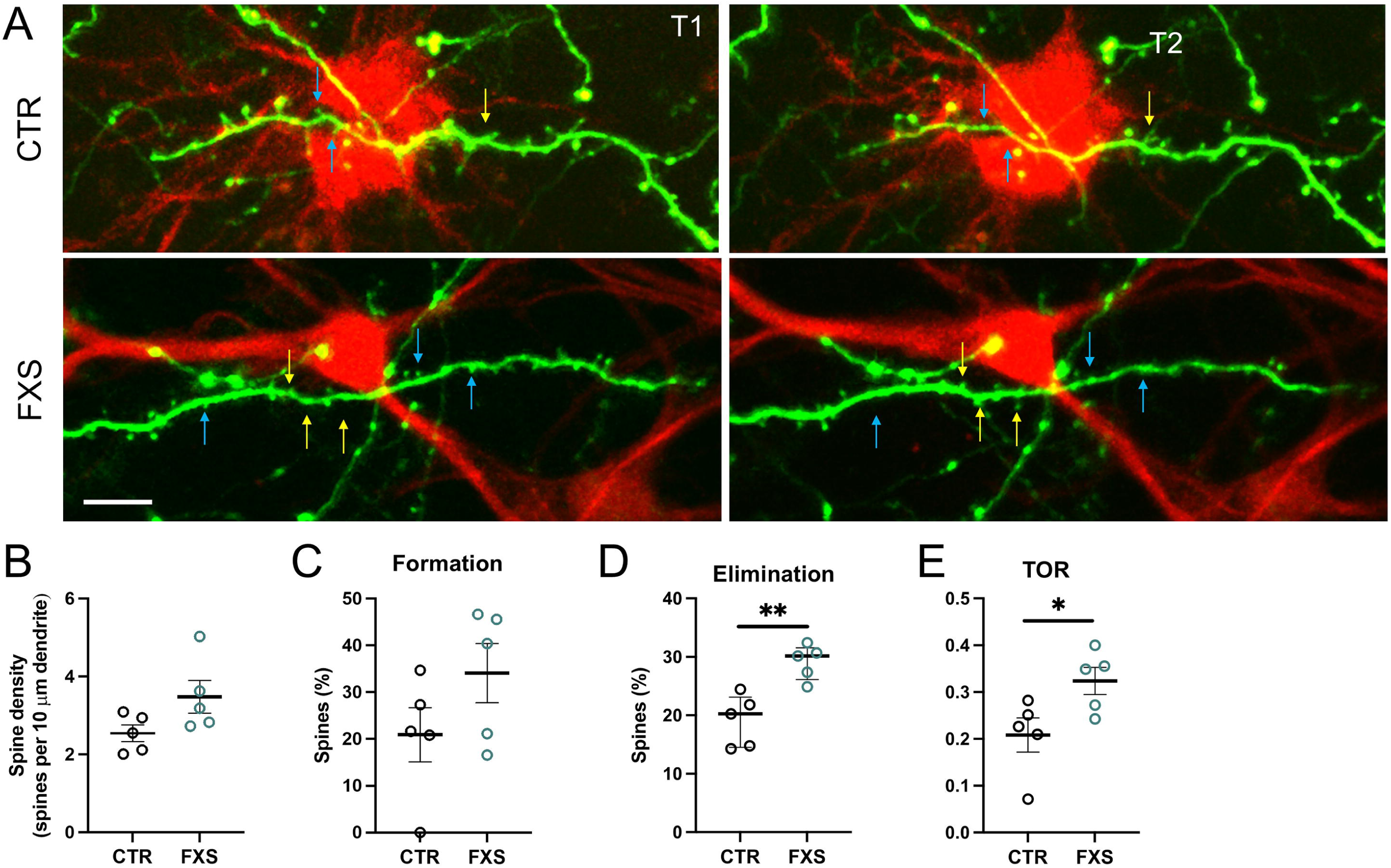
Altered dendritic spine dynamics in chimeric mice with FXS ILAs. **(A)** In vivo imaging of eGFP expressing mouse dendrites in the vicinity of CTR (top) and FXS (bottom) ILAs (distance < 20 μm on the x and y axis) in the cortex of 4-month-old chimeric mice. Repeated in vivo imaging was performed at two timepoints over a 4-day interval. Blue and yellow arrows point to eliminated and newly formed spines respectively. Scale Bar 10 μm (**B)** No differences were found in the dendritic spine density. Mann-Whitney test, P = 0.082. (**C)** No differences were found in dendritic spine formation. Mann-Whitney test, P = 0.16. **(D)** Increased spine elimination was observed in the dendrites in the vicinity of FXS ILAs. Unpaired t-test, P = 0.003. **(E)** The turnover rates (TOR) were elevated in the dendrites in the vicinity of FXS ILAs. Mann-Whitney, P = 0.037, 16-28 regions from n=5 mice in each group.

## 3. Discussion

We used hiPSCs-astrocyte chimeric mice to characterize the Ca^2+^ signaling properties of ILAs and show Ca^2+^ increases in both the cell bodies and processes. We also for the first time report the capacity of engrafted ILA processes to respond to both purinergic and noradrenergic stimulation. Although we observed no gross alterations in the development of ILA processes in FXS, ILAs exhibited enhanced ATP and NE evoked Ca^2+^ signaling. Finally, using in vivo two-photon microscopy we show increased spine elimination and turnover rates in the host dendrites in the vicinity of FXS ILAs. We conclude that Ca^2+^ signaling is altered in FXS ILAs and that the increase in the host dendritic spine plasticity is attributed to the FXS ILAs indicating the role of astrocytes in the spine dynamics in FXS.

Studies using human brain tissue and human glial chimeric mice have shown differences in Ca^2+^ wave propagation between human and mouse astrocytes [8, 28]. The study by Oberheim and coworkers [8] showed that the cell bodies and processes of human astrocytes responded to ATP. The Ca^2+^ imaging studies were performed on slices that were bulk loaded with the Ca^2+^ indicator dye Fluo-4 AM to assess the evoked responses in human astrocytes. However, fluo-4 only allows visualization and quantification of Ca^2+^ signals in somas and proximal processes [29] and is likely to conceal the Ca^2+^ events in the ILA processes. Our study utilized the genetically encoded Ca^2+^ indicator GCaMP6f that overcomes this limitation thereby revealing the Ca^2+^ signals in the ILA processes. Our study is also the first to show Ca^2+^ signaling in the ILA processes in slices and in vivo.

Rodent astrocytes respond to extracellular ATP and sensory input via elevations of intracellular calcium concentrations [30–32]. Astrocytic metabotropic (P2Y) and ionotropic (P2X) purinoreceptors are activated by ATP [1, 33]. The transcriptomic study from Zhang et al (2016) showed that mRNA expression of P2RY1, P2RY12, and P2RY13 were present in human mature astrocytes with a higher expression of P2RY1. While P2Y1 receptors are linked to PLC/IP3/Ca^2+^ signaling cascade, P2Y12 and P2Y13 receptors are linked to G_i_ proteins and inhibit adenylyl cyclase activity [1]. Interestingly, P2Y12 receptor protein expression has been observed in ILA processes in multiple sclerosis patients [34]. Low mRNA expression of P2RX4 and P2RX7 in human mature astrocytes has also been observed [9]. It is likely that the ATP-evoked Ca^2+^ increases that we observe in the ILA somas and processes are mediated by activation of one or more of these purinoreceptor subtypes that participate in the mobilization of intracellular Ca^2+^ stores.

Astrocyte α1-adrenoreceptor are primary targets for norepinephrine that are coupled into G_q_ signaling pathways which trigger increases in Ca^2+^ in rodent astrocytes following startling stimuli or the activation of locus coeruleus (LC) neurons using electrical stimulation [30, 32, 35, 36]. The α2 and β-adrenoreceptors are coupled to G_i_ and G_s_ signaling pathways, respectively [37]. Human data indicate ADRA1A, ADRA1B, ADRB1 and ADRB2 mRNA expression in mature astrocytes with a higher expression of ADRB1 and ADRB2 [9]. The norepinephrine-evoked Ca^2+^ increases observed in the ILA somas and processes are likely to be mediated by activation of α1-adrenoreceptors.

Rodent astrocytes have been shown to elicit Ca^2+^ responses with carbachol application that activates G protein-coupled muscarinic receptors (Shelton and McCarthy, 2000). However, human data indicate almost negligible levels of muscarinic cholinergic receptors and low mRNA expression for the nicotinic cholinergic receptors CHRNA4, CHRNA5 and CHRNA7. Consistent with this, we observed no responses in either the ILA somas or processes with carbachol application indicating the absence of functional muscarinic receptors. Put together, our data show that the ILAs in the chimeric mouse have functional purinergic and adrenergic receptors. Ca^2+^ elevation and propagation along the ILA processes is of potential importance as this may facilitate long-distance coordination of intracortical communication.

Long dendritic spines with immature morphologies and higher density have been observed in the postmortem brain tissue of FXS patients [38, 39]. Dendritic spine abnormalities reported in the Fmr1 KO mouse model parallel abnormalities reported in FXS patients [40, 41]. Fmr1 KO mice have increased rates of dendritic spine formation, elimination and turnover [19–21, 42]. Astrocytes have emerged as important regulators of synapse development and have been shown to promote both synapse formation and maturation [2]. Astrocytes can influence both synaptogenesis and synapse maturation through secretion of astrocytic soluble factors and the expression of these proteins has been found to be altered in Fmr1 KO mice, possibly contributing to abnormal neuronal development and altered connectivity observed in FXS [43–46]. Astrocyte-specific loss of FMRP expression has been reported to result in increased dendritic spine dynamics. These rodent studies have contributed to increasing evidence that astrocytes play essential roles in modulating the function of neurons and neural circuits.

Human astrocytes are more complex than their mouse counterparts, and abnormalities observed in *Fmr1* KO astrocytes need to be replicated in human models. Altered structural properties of ILA has been found in other neurodevelopmental disorders. A postmortem study in children with Down Syndrome showed a reduction in the number of ILA processes [47]. Our study in the chimeric mice did not reveal alterations in ILA process length in FXS. Interestingly, dendritic spine density on the dendrites in the vicinity of the engrafted FXS human astrocytes trended higher. Increased dendritic spine elimination and turnover rate were also observed in the FXS group showing that spine dynamics in the host dendrites is altered due to the FXS human astrocytes. Our study is the first to demonstrate altered dendritic spine plasticity using the chimeric mouse model and further highlights the role of astrocytes in the spine dynamics in FXS.

## 4. Materials and Methods

### 4.1. Mice

Mice were cared in accordance with NIH guidelines for laboratory animal welfare. All protocols were approved by the University of Nebraska Medical Center (UNMC) Institutional Animal Care and Use Committee. Rag1 immunodeficient mice (B6.129S7-*Rag1^tm1Mom^*/J, Jackson Laboratory, IMSR Cat# JAX: 002216, RRID: IMSR_JAX: 002216) were bred at the UNMC facility with a 12-hr light/dark cycle with food and water available ad libitum.

### 4.2. Stem cell differentiation

The FX11-9u (RRID: CVCL_EJ77), FX11-7 (RRID: CVCL_EJ76) hiPSC lines, and WA01 (RRID: CVCL_9771), H1-FMR1-KO hESC lines were obtained from WiCell. The FX11-9u and FX11-7 hiPSC lines were derived from the same FXS patient. The FX11-9u line retained the FMRP expression due to the mosaicism of CGG repeats during reprogramming [48]. The H1-FMR1-KO (abbreviated as FMR1-KO) was engineered by CRISPR/Cas9 targeting exon 3 of the FMR1 gene in WA01 [49]. The SC176 control line was a gift from Dr. Gary Bassell [50]. The same protocol as previously described [15, 16] was used for RFP transduced cells. For the GCaMP6f and mScarlet transduced cells, the embryoid bodies were generated by treatment of hiPSCs with ReLeSR^TM^ (STEMCELL Technologies) and passing the clumps though a 100 µm sterile strainer.

### 4.3. Viral transduction

NPCs were transfected with CMV-RFP lentivirus (Cellomics Technologies) or LV-CMV-GCaMP6f-T2A-mScarlet (SignaGen Laboratories) as previously described [15] with the modification that transduced NPCs were expanded prior to differentiation to astrocytes for all cells except those transduced with CMV-RFP.

### 4.4. Engraftment of human iPSC derived astrocytes

Engraftment was performed as previously described [15]. Briefly, Rag1^-/-^ neonatal mice (both males and females) were transplanted on postnatal day 1 with hiPSC-derived astrocytes expressing GCaMP6f and mScarlet or RFP in a random manner. The pups were cryoanesthetized for 4 min and transferred to a neonatal stage (Stoelting) that was cooled to 4°C during the stereotaxic injections. For cortical labeling, the pups were injected directly through the skin into two sites: AP −1.0 and −2.0, ML ± 1.0 mm, ventral 0.2-0.8 mm, 10,000 cells/µl per site using a Hamilton syringe.

### 4.5. Brain slice preparation

At 6 months following hiPSCs-Astrocyte (hi-Astrocyte) engraftment, anesthetized mice (Avertin, 0.25 mg/g body weight) were transcardially perfused with carbogenated (95% O_2_/5% CO_2_) N-Methyl-D-glucamine (NMDG) artificial cerebrospinal fluid (aCSF) containing (in mM) 92 NMDG, 2.5 KCl, 1.25 NaH_2_PO_4_, 30 NaHCO_3_, 20 HEPES, 25 glucose, 2 thiourea, 5 Na-ascorbate, 3 Na-pyruvate, 0.5 CaCl_2_ and 10 MgSO_4_ [51]. Following perfusion, mice were decapitated and their brains quickly removed and immersed in ice-cold carbogenated aCSF. Acute coronal slices containing the frontal cortex were cut to 300 µm using a vibratome. The slices were transferred into a pre-warmed chamber containing carbogenated NMDG aCSF and held for 10 minutes at 32–34 °C. After this initial recovery period, the slices were transferred into a new holding chamber containing room-temperature recording aCSF containing (in mM) 119 NaCl, 2.5 KCl, 1.25 NaH_2_PO_4_, 24 NaHCO_3_, 12.5 glucose, 2 CaCl_2_ and 2 MgSO_4_ under constant carbogenation and held in this chamber for at least 1 hour before the imaging commenced. For imaging, slices were transferred to the submersion-type chamber and superfused at room temperature (22–24 °C) with recording aCSF saturated with 95% O_2_/5% CO_2_.

For the Ca^2+^ imaging data shown for 4 months following hi-Astrocyte engraftment, anesthetized mice were transcardially perfused with carbogenated aCSF. Acute coronal slices were cut in carbogenated aCSF containing (in mM) 126 NaCl, 3 KCl, 1.25 NaH_2_PO_4_, 4 MgSO_4_, 2 CaCl_2_, 26 NaHCO_3_, and 10 dextrose. The slices were transferred into a holding chamber containing room-temperature recording aCSF containing (in mM) 126 NaCl, 3 KCl, 1.25 NaH_2_PO_4_, 1 MgSO_4_, 2 CaCl_2_, 26 NaHCO_3_, and 10 dextrose.

### 4.6. Dye loading in slices

Following a recovery period, slices were loaded with Fluo-4 AM (5 µM) and 0.6 µL of Pluronic acid F-127 diluted in 1.5ml of aCSF (containing 1 mM Mg^2+^) saturated with 95% O_2_/5% CO_2_ for 40 minutes at room temperature. Following three 10 minutes washes with aCSF, the slices for imaging were transferred to the submersion-type recording chamber and superfused at room temperature (22–24°C) with aCSF saturated with 95% O_2_/5% CO_2_ containing.

### 4.7. Viral injection and cranial window implantation

To perform *in vivo* multiphoton imaging of eGFP expressing mouse neurons in the vicinity of RFP-expressing human astrocytes, we performed viral injection of AAV1.CaMKII.0.4.Cre (1:5,000) and AAV1.CAG.FLEX.eGFP into Layer 2/3 of the cortex and implanted a cranial window in 3.5 -month-old chimeric mice [26]. To visualize Ca^2+^ activity in vivo, a cranial window was implanted in 6-month-old chimeric mice expressing GCaMP6f and mScarlet. Mice were injected twice daily with 5 mg/kg enrofloxacin for 6 days and 5 mg/kg carprofen daily for 20 days. Mice were allowed three weeks to recover from the surgery prior to imaging.

### 4.8. Multiphoton imaging

#### 4.8.1. Slice imaging and pharmacology

Time-lapse imaging was performed with a two-photon microscope (Moving Objective Microscope (MOM), Sutter) attached to a Ti : sapphire laser (Chameleon Vision II, Coherent) using a 25x water immersion objective (1.05 NA, Nikon) and equipped with a resonant scanner and a piezo stage (nPFocus400, nPoint). Excitation wavelength was tuned to 920 nm with 10-12 mW power as measured at the back aperture. Two-channel detection of emission wavelength was achieved by using a 670 nm dichroic mirror and two external photomultiplier tubes (GaAsP). A 535/50 bandpass filter was used to detect GCaMP6f emission wavelength, and a 610/75 bandpass filter was used to detect mScarlet or RFP. For imaging, we used ScanImage software (Vidrio) written in MATLAB v. R2024b (The MathWorks) [52]. Images of ILA were acquired at a resolution of (0.3 µm/pixel), a 20 µm volume at a step size of 1 µm and a frame rate of 1 Hz. Time-lapse imaging was performed for a period of 5 minutes. For each slice, two minutes of spontaneous Ca^2+^ activity was recorded first, followed by agonist application for 2 minutes. Agonists were added to the perfusing ACSF at the following final concentrations: 100 µM ATP, 50 µM norepinephrine (NE) and 50 µM carbachol. Slices were exposed only to a single agonist.

#### 4.8.2. *In vivo* imaging

##### Dendritic spine imaging

On the imaging day, four-month-old chimeric mice were anaesthetized with a ketamine/dexdormitor mixture (100 mg/mL and 0.5 mg/mL, respectively, dosage 2.5mL/kg). Excitation power measured at the back aperture of the objective was typically around 20 mW and was adjusted to achieve near identical levels of fluorescence for each imaged region. Each optical section was collected at 512 x 512 pixels (0.186 µm/pixel). During an imaging session, 5 to 10 ROIs per animal were selected along with the apical dendritic tufts of eGFP expressing neurons within the vicinity of the RFP expressing human astrocytes in the cortex. Each ROI consisted of a stack of images (50-80 optical sections, separated axially by 1 µm). The coordinates of each ROI were recorded using the XYZ motor on the MOM for subsequent imaging days. After imaging, mice were revived from anesthesia with antisedan (atipamezole hydrochloride 5.0 mg/mL). Dendritic spines were imaged twice, at a four-day interval.

##### In vivo calcium imaging

Mobile home cage (MHC, Neurotar) was used for imaging in awake mice where a head-fixed mouse can move around in an air-lifted MHC that features a flat floor and tangible walls and explore the environment under stress-free conditions. Prior to the imaging experiment, the mouse was habituated to the MHC by gradually increasing the duration of the habituation sessions every day and acclimating the mouse to the sounds of the laser scanning mirrors [26]. The habituation phase was started two weeks after the cranial window was implanted. Time-lapse imaging was performed every 2s for a period of 5 minutes. Each optical section was collected at 512 × 512 pixels, 0.37 μm/pixel and a 20 µm volume was acquired at a step size of 1 µm.

### 4.9. Image Analysis

Process length analysis was performed as previously described [15]. Briefly, in each section multiple 5-pixel broad straight lines, 150–200µm apart, were drawn from the pial surface to the deeper layer. For each line, the plot profile function was used to estimate the pixel intensity values along the line. The location of the last peak of RFP fluorescence was used as a measure of the extent to which processes traversed across the cortex and averaged per section. The values from control FX11-9u line were previously published [15]. Analysis was performed while blinded to engrafted line identity.

#### 4.9.1. Analysis of astrocytic Ca^2+^ events

The images acquired as a 20-micron volume were maximum intensity z-projected in 5 micron volumes for further analysis. For proportion of active ILA processes up to 10 processes per field of view were scored as to whether they were responsive to an agonist. Event-based analysis of astrocyte Ca^2+^ image events was performed using Astrocyte Quantitative Analysis (AQuA 2020) software [53] only on a subset of well isolated processes (1-3 per field of view). The AQuA-detected events were categorized based on area of the Ca^2+^ events with 3< 10 µm^2^ (small) and >10 µm^2^ (large). Not all processes displayed both small and large sized events. For analysis of astrocytic Ca^2+^ activity in the soma, a region of interest was outlined around the soma. The Ca^2+^ events were categorized into spontaneous and agonist-evoked events. The peak amplitude (dF/F), area under the curve, duration (at half width), rise time (10%-90%) and decay time (90%-10%) of the events were analyzed. Clampfit v10.6 (pCLAMP, Molecular Devices) software was used to detect and measure the parameters for the soma. Analysis was performed while blinded to engrafted line identity.

#### 4.9.2. Analysis of spine plasticity

The analysis of spine plasticity was performed on ImageJ software. For each image, dendrites with at least 25 µm-length, as well as located within 5 focal planes and 20 µm XY coordinates from the RFP-expressing astrocytes were selected for further analysis. All spines on the selected dendrites were counted and tracked over time to identify newly formed, eliminated and stabilized spines. Dendritic filopodia were distinguished as long dendritic protrusions with no head and were excluded from analysis (< 3% in both genotypes). Dendritic spines were analyzed by scrolling through individual z-planes within a stack. Spines were categorized as stable if they were present in the previous image as well as the one being analyzed, eliminated if they appeared in the previous image but not in the image being analyzed, newly formed as they appeared in the image being analyzed but not in the previous image. The percentage of spine formation and elimination was calculated using the number of newly formed or eliminated spines to the total number of spines analyzed in the image, respectively. Turnover rate (TOR) was calculated as the ratio of the sum of newly formed and eliminated spines to twice the total number of spines at baseline [20]. Analysis was performed on raw unprocessed images. For presentation purposes, images were despeckled and went through maximum intensity projection of 3–15 planes of focus. Analysis was performed while blinded to engrafted line identity.

### 4.10. Tissue preparation and confocal imaging

Mice were deeply anesthetized with TribromoEthanol (Avertin, 400mg/kg i.p.) and transcardially perfused with 4% paraformaldehyde in phosphate buffer (0.1 M) at 3 and 9 months post-engraftment. The brain was dissected, post-fixed overnight, and 100 µm sagittal sections were cut on a vibratome in PBS. Confocal imaging of tissue sections was performed on a Nikon A1R upright microscope and images were acquired using a 20x (0.75 NA) objective. Images were collected at 512 × 512 pixels (with a pixel size of 1.24 μm) and a step size of 1 μm with 561 nm laser.

### 4.11. Statistical analysis

Data were analyzed using GraphPad Prism. Outliers were removed using the ROUT approach (Q=1%). Normal distribution was tested using the Shapiro-Wilk test. Data are reported as mean ± S.E.M or median ± interquartile range. On scatter plots, medians are denoted by a thin line and means by a thick line. The following tests were conducted as appropriated: unpaired t-test, Mann-Whitney test and two-way ANOVA. P values was adjusted for multiple comparisons. For proportion of responding ILA processes, a Chi-Square test was used.

## Supplementary Materials

The following supporting information can be downloaded at: https://www.mdpi.com/article/doi/s1, Figure S1: Enhanced ATP-evoked Ca^2+^ signaling in FXS astrocytes at 4 months; Video S1: ATP-evoked Ca^2+^ signaling of ILAs in slices. Video S2: Ca^2+^ signaling of ILAs imaged in vivo through a cranial window.

## Institutional Review Board Statement

All protocols were approved by the University of Nebraska Medical Center (UNMC) Institutional Animal Care and Use Committee.

---

**Supplemental Figure 1: Enhanced ATP-evoked Ca^2+^ signaling in FXS astrocytes at 4 months**

**(A)** Time-lapse imaging of CTR and FXS astrocytes in a cortical slice from a 4-month-old chimeric mouse. Images of RFP-expressing astrocytes (red) and time-lapse images of the fluo-4 channel (green) showing ATP-evoked Ca^2+^ responses (top panel: CTR, bottom panel: FXS). Traces for the Ca^2+^ responses are shown for the regions of interest (ROI 1 and 2) indicated in the RFP images. **(B)** Peak amplitude of ATP-evoked Ca^2+^ responses is higher in FXS astrocytes. Mann-Whitney test. P = 0.0023. Frequency of ATP-evoked Ca^2+^ responses. Mann-Whitney test. P = 0.2699. Duration of ATP-evoked Ca^2+^ responses. Mann-Whitney test. P = 0.1807. N = 6-8 slices from 4-5 mice in each group.

