## Supplementary figures and images for "Activity of human-specific Interlaminar Astrocytes in a Chimeric Mouse Model of Fragile X Syndrome"

### Supp Fig.1 Ca imaging slices ILA soma.tif

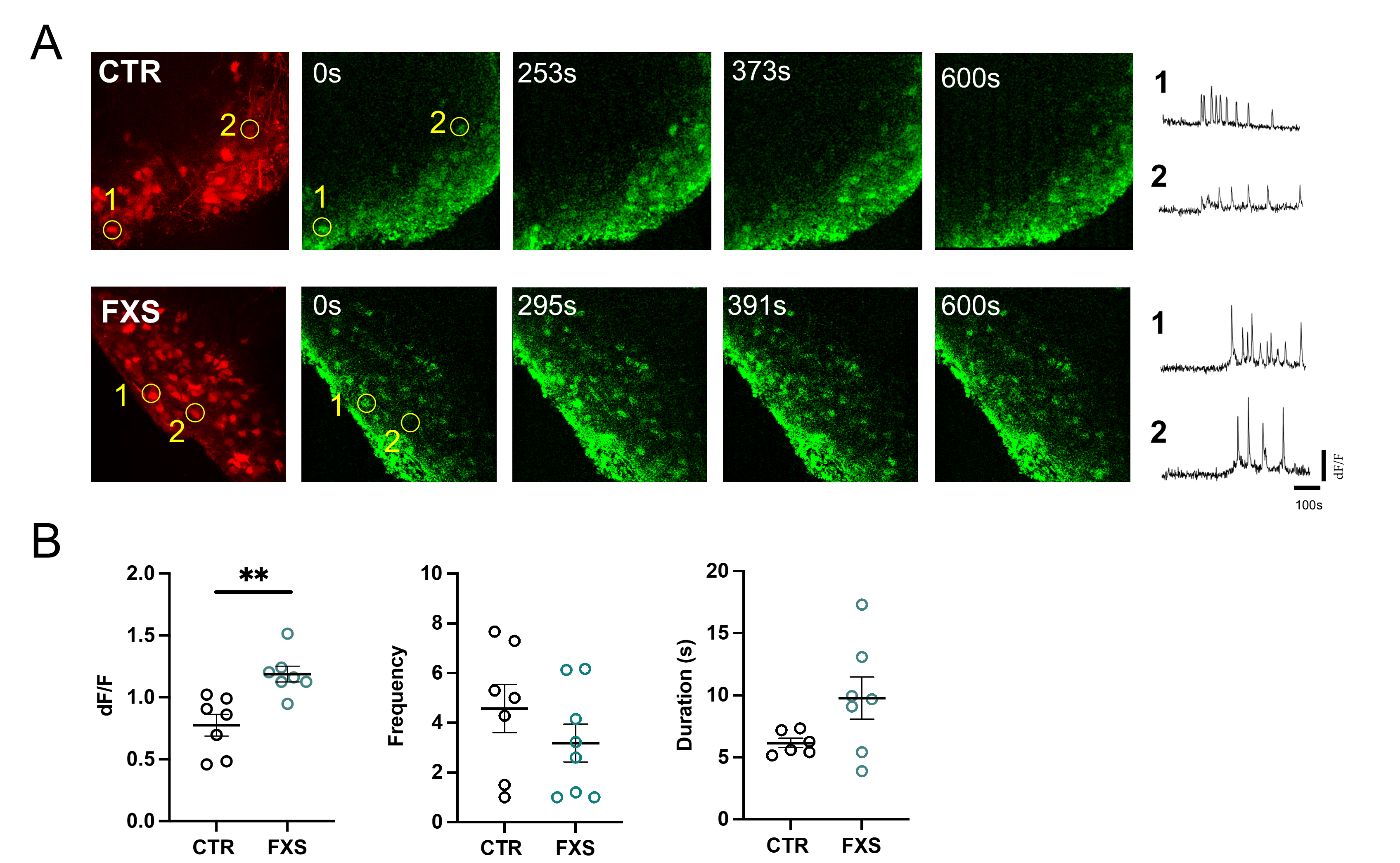
